# Alanine-scanning mutagenesis library of MreB reveals distinct roles for regulating cell shape and viability

**DOI:** 10.1101/2024.04.02.587816

**Authors:** Suman Maharjan, Ryan Sloan, Jada Lusk, Rose Bevienguevarr, Jacob Surber, Randy M. Morgenstein

## Abstract

The bacterial actin-homolog MreB is a crucial component of the rod-system (elongasome) that maintains rod shape in many bacteria. It is localized beneath the inner membrane where it organizes the elongasome complex. Depletion or deletion of *mreB* results in loss of rod shape and cell death; however, the mechanism of how MreB operates is not known, given that the protein cannot be purified in a functionally intact form. Past studies have reported mutations in *mreB* cause varying degrees of cell shape and size alterations based on the type and position of the substitution. To better understand the role of MreB in rod shape formation we have taken the first truly systematic approach by replacing the native copy of *mreB* with an alanine-scanning mutagenesis library. Surprisingly, we observed stably growing spherical mutants that have lost MreB’s function(s) for shape regulation without losing viability. Hence, MreB has vital functions related to growth in addition to shape maintenance that can be separated. In support of this, rod shape suppressor analysis of these spherical mutants only revealed reversions or intragenic *mreB* mutations, suggesting that MreB is indispensable for rod shape. Additionally, our results imply the elongasome is no longer active in these strains, suggesting a novel way for rod shaped bacteria to synthesize cell wall.

## Introduction

The cell shape of bacteria is determined by the extracellular peptidoglycan (PG) cell wall and is ^1–8^ tightly maintained to ensure efficient growth, division, and survival^1,9,10^. Loss of PG in rod bacteria results in the formation of osmosensitive spherical L-forms in which rod shape can be regained upon new cell wall synthesis in an MreB dependent manner ^2,7,11–14^. Most rod shaped bacteria maintain their shape due to the activity of a complex of proteins termed the ‘elongasome’ or ‘rod system’^10,15,16^. The elongasome consists of a partner SEDS (Shape, elongation, division and sporulation) family transglycosylase (RodA) and penicillin binding protein (PBP) transpeptidase (PBP2), in addition to the bifunctional enzyme PBP1A to promote cylindrical growth in addition to MreCD ^17,18^. Although their function in unknown, it has been suggested that MreC regulates the elongasome through PBP2 and RodA activation while MreD may suppress such interactions^19–23^. These enzymes are regulated by MreB whose localization and polymerization are affected by the transmembrane protein RodZ^24–27^. MreB is biologically active as cell membrane associated polymeric antiparallel double nanofilaments in the cytoplasm^25,28–32^. Rod-shaped bacteria may carry one (*E. coli, Enterobacteriaceae, Caulobacter crescentus, Anabaena, Nostoc*) or two or multiple paralogs (*Thermotoga maritima*, *Bacillus*, *Spiroplasma*) of MreB ^33–42^. In *E. coli,* MreB is localized to the central cylindrical region of the cell in multiple discrete patches correlating to the location of new PG synthesis^24,25,30,32,43–48^. The exact mechanism of how MreB operates is not completely understood.

As the main protein for rod shape maintenance MreB is not only essential for rod shape but also growth. In *B. subtilis*, which has three paralogs *mreB*, *mbl* and *mbH*, the triple deletion mutation is lethal while loss of any one paralog results in distorted shape^37^. In *C. crescentus*, depletion of *mreB* leads to lemon-shaped cells while deletion of *mreB* is lethal^34,36,49^. In *E. coli*, depletion or deletion of *mreB,* or any other *mre* proteins, leads to loss of rod shape and eventually cell death^50^. While MreB appears to be essential for viability it can be suppressed through very slow growth or the overexpression of the division proteins, FtsZAQ, or their regulator SdiA, suggesting that the original discovery of these genes was in a strain with increased levels of FtsZAQ^51–54^. While these suppressors allow cells to grow without *mreB* they do not restore shape, suggesting the MreB is essential for rod shape^1,50,51,55^.

While it has been established that MreB is necessary for rod shape, how MreB achieves this is not known. Past studies suggest MreB acts upstream of other shape determinants, as point-mutations can modulate one or more morphological parameters to create different rod-like shapes^56^. Early studies focused on the conserved motifs of the Actin/Hsp70 superfamily in different bacterial MreBs. In *B. subtilis*, introduction of a mutation in the phosphate 2 motif of MreB that corresponds to the lowering of ATPase activity of eukaryotic actin leads to longer MreB filaments resulting in shape and growth defects and the mislocalization of other *mreB* isoforms^57^. In *C. crescentus*, cells were selected for A22 resistance and screened for cell shape defects. Mutations in *mreB* that modulate cell shape were enriched for in the nucleotide-binding pocket of MreB at the same conserved motifs as tested in *B. subtilis*^58,59^. While in *E. coli,* A22 has been used to find *mreB* mutations that alter cell width^46^. Long term evolution experiments also identified MreB mutations resulting in altered morphology and changes in growth rate^60–62^. The most comprehensive study of MreB involved FACS-sorting cells for shape defects from a plasmid complementation library of *mreB* mutants generated through error-prone PCR mutagenesis. They identified a number of point mutations modulating cell size without affecting growth rate in *E. coli*^56^. In some instances different morphologies are obtained when different amino acids are substituted at the same position^27,33,46,63^. Together these observations suggest amino acid residue and position can independently tune rod-shape.

Each of these prior experiments has drawbacks. The use of A22 to isolate mutants is inherently biased as they most likely will not select for mutations across the entire MreB protein and will miss any residues important for shape regulation independent of A22 resistance. The mutations picked from evolution experiments are confounded by the fact that these strains produced multiple mutations in cell shape determinants, not just *mreB* mutations, and may be specific to the media in which the cells were grown. While complementation of an *mreB* deletion with a mutant plasmid library allows for an unbiased approach, it possibly introduces copy number effects from the plasmid as well as the possible loss of regulatory sequences. In addition, there may be bias in the possible shapes able to be selected for by cell sorting. The studies on bacteria with multiple MreB isoforms have functional redundancy making the relevancy of homology-based prediction into *E. coli* MreB (*Ec*MreB) unreliable.

The lack of efficient protocols to purify functional mono- or poly-meric MreB from *E. coli* has hindered any physical and biochemical interrogations, making *in-vivo* genetic experiments even more important. Hence, this necessitates more work beyond homology-based extrapolation to understand the role of the residue(s) involved in shape maintenance and/or determination. To overcome some of these previous biases, we have taken a systematic approach to study *Ec*MreB by creating a fully functional GFP-tagged native-site alanine-scanning mutagenesis library to assess the impact of each residue on cell shape determination. Our construct is chromosomally located at the native site making it the sole copy of MreB under its native promoter and regulatory elements. Here, we present a detailed shape analysis of the 337-point mutants in this library with an emphasis on 27 spherical mutants. Our results suggest that MreB has independent functions for rod shape maintenance and growth. A rod shape suppressor screen supports the idea that MreB is absolutely necessary for rod shape. These spherical mutants are resistant to the PBP2 targeting antibiotic mecillinam and A22 suggesting they no longer rely on the elongasome for PG synthesis suggesting a possible novel cell wall synthesis mechanism.

## RESULTS

### Most *mreB* alanine substitutions are viable

MreB is an important protein in the regulation of cell shape. While multiple studies have been performed to understand how MreB functions, we still have little idea of how MreB regulates cell shape and the role of individual residues. Past experiments have used antibiotics or plasmid-based systems to make random MreB point mutations^28,29,34,46,47,50,50,55,56,58,59,63–65^. These experiments have shown that different changes at a single amino acid site can tune cell size; however, there has been a lack of a systematic approach to understanding the role of each amino acid residue in cell shape regulation.

To systematically understand how MreB functions, we constructed a native-site GFP-tagged alanine-scanning mutagenesis library of *mreB* via lambda red recombination. All amino acids were individually changed to an alanine except for the native alanines, which were changed to leucine, and the initiating methionine, which was left unchanged. The construct used to make the gene replacements has *msf*GFP fused to *mreB*^46^ in an internal non-conserved loop (sandwich fusion) with flanking regions of homology on both sides of *mreB* and a kanamycin resistance cassette (**Fig. S1A**). Cells expressing the lambda red plasmid PKD46 were transformed with the constructs, selected on kanamycin LB agar plates, and screened for GFP fluorescence. The strains with mutations downstream of the GFP were sequenced to ensure the desired recombination occurred. Out of the 346 amino acids of MreB, we were unable to make nine amino acid substitutions (**Table S1, Fig. S1C**) leaving us with a library of 337 mutants. Seven of these mutations do not occur in suspected self-interaction regions of MreB which were thought to be important for MreB assembly and function. Our inability to successfully transform strains with the mutant *mreB* alleles suggests these residues are important for viability. In support of this, these residues are highly conserved among both Gram-negative and Gram-positive species (**Table S1**).

### *E. coli* can stably grow as spheres

To determine the role of each residue of *mreB* on cell shape we imaged the entire library grown in LB at 37°C. Due to the large number of strains that were imaged we arrayed the mutants in 96-well plates with a wild-type (WT) strain in each plate and fixed the cells before imaging. A Δ*rodZ* mutant was included as a control for a non-rod shape phenotype. To keep all cells in exponential phase we serially diluted the cultures as previously done (see materials and methods). Cells were fixed in paraformaldehyde to preserve their shape and fluorescence during the imaging process. Surprisingly, this initial screening revealed a wide variety of cell shape phenotypes, with some mutants having disrupted shapes similar to Δ*rodZ* cells.

Because MreB is known to be involved in rod shape determination we expected many mutations to affect rod shape. However, based on past experiments we did not expect to find a complete loss of rod shape as this was expected to be lethal. To quantify how rod-like cells are, we measured the coefficient of variation of the intracellular diameter deviation (IDD) of all cells in the library. This metric measures the standard deviation of the cell diameter across the long access of the cell^25^. The more spherical a cell is the larger the IDD, as a rod will only have changes to its diameter at the poles. Due to the number of mutants observed we grouped cells into a histogram based on IDD values. Most of the MreB mutants have either no change in rod shape or a distorted rod-like shape phenotype (left side of the histogram); however, we observed mutants with shape defects beyond the rod-like spectrum (**Fig. 1**). The cells that have the least rod-like shape (the last four bins in **Fig. 1A**) were reimaged using live cell imaging. As the length of a rod will determine what percent of the cell is a pole, thereby effecting the IDD value (**Fig. S2A**); therefore, we measured the aspect ratio of the cells in last four bins from the IDD histogram and found that the average aspect ratio for each bin was 3.03 ± 0.43, 2.70 ± 0.47, 2.67 ± 0.45 and 2.14 ± 0.12 respectively. Using an aspect ratio of 3 as a cutoff for rod-shape, the last three bins will be considered non-rod-like and excluded from future cell shape analysis. Interestingly the Δ*rodZ* mutant has the highest IDD value in the penultimate bin indicating that the strains in the black bin have completely lost rod shape.

**Figure 1.**
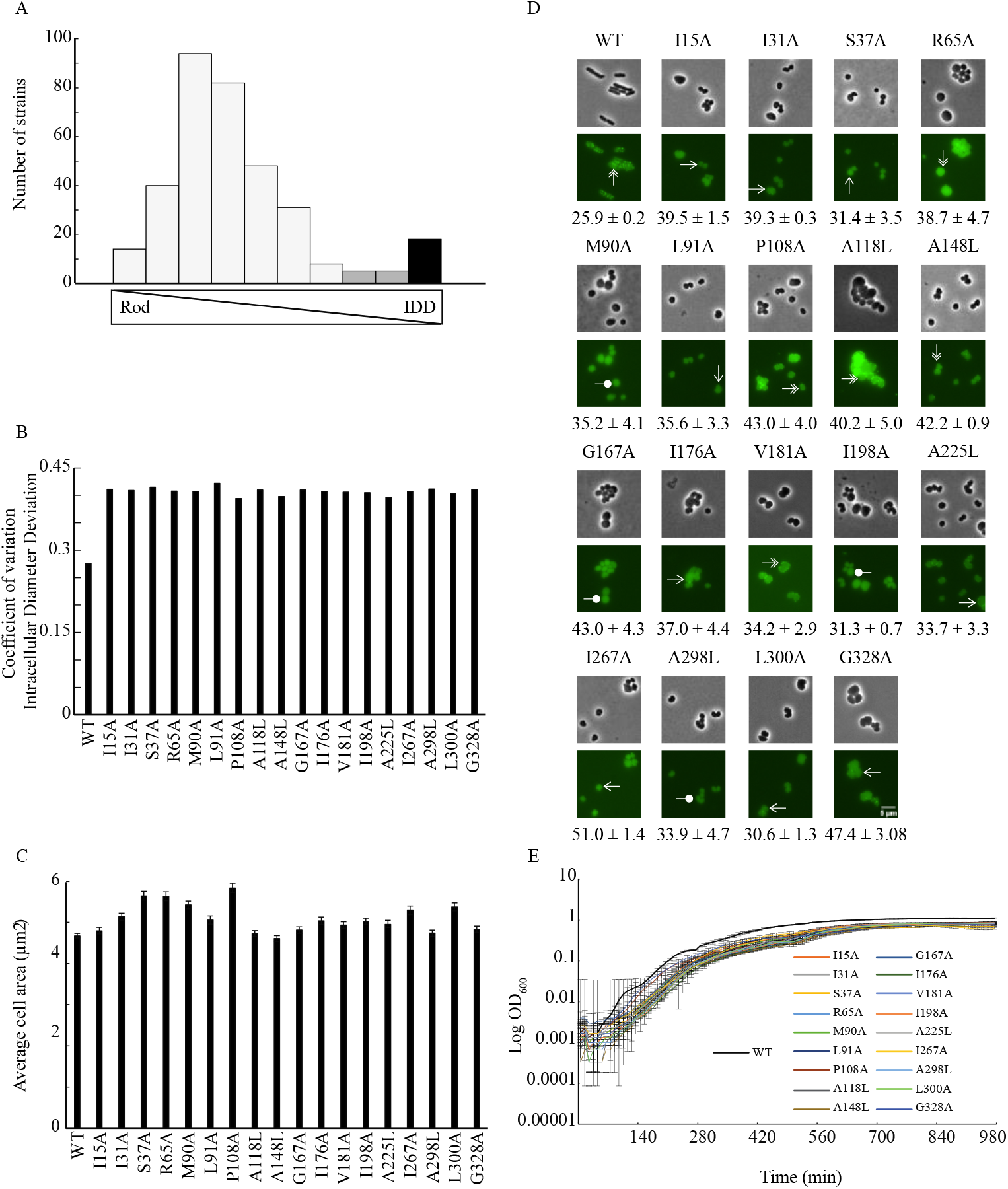
Alanine-scanning mutagenesis library shows non-rod shape phenotypes. A) Histogram of 337 point-mutants (n >100 cells for each strain) based on the coefficient of variation of intracellular diameter deviation (IDD). Black bar indicates the 18 mutants with the highest IDD, while gray bars indicate the mutants considered to have lost rod shape. Cells were grown in 96-well plates as described in materials and methods and then fixed. B-C) Live cell imaging and quantification of WT and the eighteen mutants (n >600 cells for each strain) from the black bin with extreme shape defects. Cells were grown in LB at 37°C to early exponential phase. Data is pooled from three independent experiments. B) IDD measurements and C) mean cell area. D) Representative phase contrast and fluorescence images of the eighteen mutants and WT grown in LB at 37°C to an OD 0.2-0.25. The numbers represent max doubling times in min (mean ± sd). Double arrow heads point to the cells with WT-like MreB foci, single arrow heads indicate cells with mostly cytoplasmic MreB but some foci, and circle heads point to cells with cytoplasmic MreB. E) Representative growth curves of eighteen round mutants and WT. Cells were grown overnight in LB and sub-cultured in equal numbers into a 96-well plate with three replicates for each strain. Error bars are the standard deviations from three wells. The black line represents the growth curve for WT.

### Spherical *mreB* point mutant do not have known suppressor mutations

Because it was unexpected to be able to stably grow spherical *E. coli* with a broken MreB, we performed whole genome sequencing (WGS) to check for the presence of known suppressor mutations. We chose ten of the spherical mutants for WGS and found no mutations in the PG synthesis machinery (elongasome and divisome components), their known regulators, nor their promoter regions. However, we did find that our parental MG1655 WT strain used to make the library contain a point mutation in the bifunctional ppGpp hydrolase/synthetase, *spoT* (*spoT_A26E_*), while the WT fluorescent strain (NO50) does not (**Tables S2-3**). The levels of ppGpp have been shown to regulate cell size and may be linked to mecillinam resistance in *E. coli*^66,67^. SpoT is the major hydrolase for ppGpp with weak synthetase activity^68,69^. While this mutation is outside of the known synthetase activity region (amino acids 67-374) it falls within the hydrolase region and may be affecting intracellular levels of ppGpp and helping to stabilize mutants with a broken MreB^70^. To test if this mutation aids in the viability of our round mutants, we moved the WT *mreB-GFP_sw_* fusion into the parental MG1655 WT strain (*spoT_A26E_*) and compared its susceptibility to A22 and mecillinam with that of the MG1655 (*spoT_A26E_*) and our fluorescent NO50 parent (*spoT_WT_*). If this *spotT* mutation was changing ppGpp levels, we would expect to see increased mecillinam resistance. We found that the *spoT_A26E_* mutation does not lead to A22 or mecillinam resistance suggesting that the *spoT_A26E_* mutation does not provide any advantage for viability and growth for the mutants in this study. Hence, the growth of these spherical mutants suggests MreB may be playing a key role in viability that is separate from its role in cell shape determination.

### Spherical strains have a variety of MreB localization patterns

A point mutation in *mreB* could lead to loss of rod shape due to lower levels of MreB, loss of interaction with an elongasome component, loss of self-interactions, changes to polymer characteristics, or changes to ATP binding/hydrolysis. We observed that the round mutants have both different MreB localization patterns and fluorescence intensities. A loss of monomer-monomer interactions should result in cytoplasmic localization of MreB, which is seen in four mutants (**Fig. 1D**, circle head), although none have mutations in the predicted monomer-monomer interface^71^. Similarly, we would expect mutations that block protofilament interactions to have most of the MreB in the cytoplasm. Again, we see multiple mutants with either only cytoplasmic or mostly cytoplasmic MreB localization (few noticeable foci) without having mutations in the predicted protofilament interface (single arrowhead)^28^. WT MreB forms multiple foci across the cell body. This phenotype is seen in multiple mutants suggesting that polymerization is not broken in these strains (double arrowhead), of which two, R65A and V181A lie in the predicted monomer-monomer interface. The presence of foci suggests the MreB in these mutants has lost the ability to interact with an essential partner resulting in filaments (foci) but not rod shape or that these mutations have altered the physical characteristics of the polymers so that they or the elongasome complexes are no longer functional. Lastly, we noticed differences in MreB concentration across the different strains as seen by the fluorescence intensity (**Fig. 1D**). This further supports MreB having distinct roles for shape regulation and viability as mutants that express lower levels of MreB may not have enough MreB to regulate rod shape but must have a sufficient MreB concentration to support growth.

### Residues involved in shape and viability are highly conserved

After live cell imaging, strains with the most extreme shape defects (dark bins) all appear misshapen with a high IDD value (**Fig. 1B, S2B**). We quantified the growth rate of these mutants in LB at 37°C and found that all strains from these bins have a longer doubling-time than WT cells (**Fig. 1C-D, S2C-D**). Interestingly, while these strains all have high IDD values, they have different cell sizes (**Fig. 1E, S2E**) suggesting that MreB is being disrupted in separate ways leading to different types of spherical cells. Additionally, MreB localization in the cells with the highest IDD is different, displaying both diffuse signal and discrete aggregates or foci (**Fig. 1D**). We compared the conservation of the residues that lead to the 27 non rod-shaped mutants in a variety of Gram-negative and positive organisms. 69.4% of the residues that result in nonviable or misshapen cells (25/36) are identical among the tested species compared to only 36.3% of the total protein (**Table S1**). The increased conservation of these specific residues over the total protein further supports the conclusion that they are important for MreB functions.

### Cell width is tightly controlled by MreB

MreB has been linked to cell width or rod diameter regulation in *C. crescentus* and *E. coli*^46,47,72,73^. To determine which residues are important for width regulation we measured the cell width of all the mutants (Supp Data). Due to plate-to-plate variances in cell growth and size, we calculated the foldchange of the cell width of each mutant to the WT strain in each individual plate and represented the data as a histogram (**Fig. 2A**). 60 mutants represented by the black bar (**Fig. 2A**) have a low fold change indicating these mutants are thinner than WT. We performed live cell imaging on the 20 mutants with the smallest ratio after growth to early exponential phase (OD 0.2-0.25). 19 (95%) were significantly thinner than WT cells and have a decrease of cell width by 6.10% to 22.54% (*p* <0.0001) with respect to WT cells **(Fig. 2B**) indicating that our initial fixed-cell screen accurately measures thin cells. Previous results found a weak correlation between the width of cells and the coefficient of variation of width; therefore, we measured the coefficient of variation of width (CV_width_) of these 19 thin mutants and found that almost all of them have a smaller coefficient than WT cells **(Fig. 2C**) suggesting that as cells get thinner, they have a more consistent width across the population^56^.

**Figure 2.**
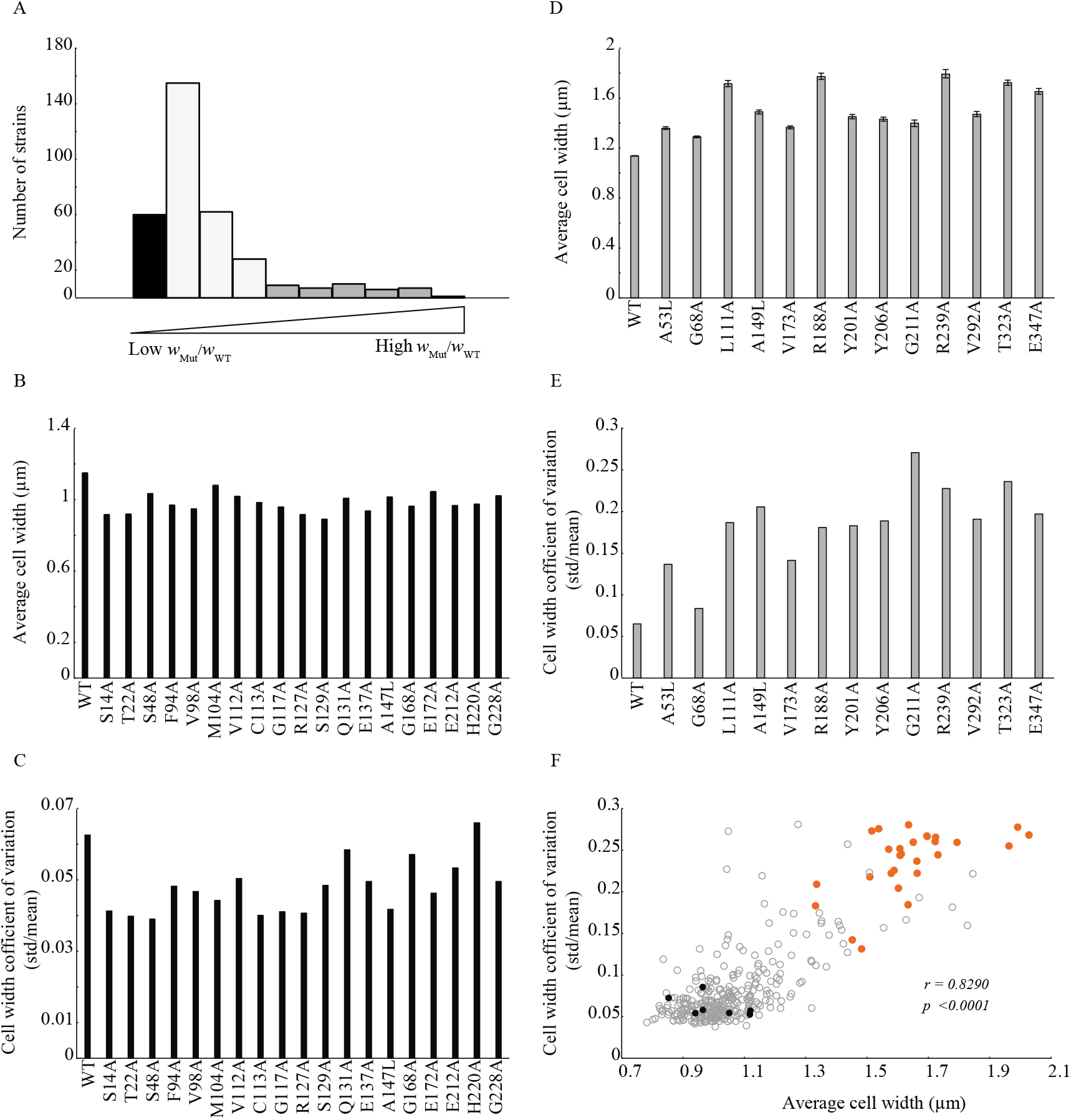
MreB robustly controls cell width. A) Histogram representing the cell width foldchange of 337 point-mutants (n >100 cells for each strain) compared to WT from fixed cell imagin. Cells were grown in multiple 96-well plates, with a WT in each plate. The black bar represents the thinnest mutants, and the gray bars represent the widest mutants. B-E) Live cell imaging of cells grown in LB at 37°C to an OD 0.2-0.25. B) Average cell width of the twenty thinnest mutants (black bar from A) and WT cells (n >400 cells for each strain, *p <0.0001*). Error bars are 95% CI. C) Coefficient of variation of cell width among the thin mutants and WT. D) Average cell width of wider mutants (gray bars from A) and WT (n >600 cells for each strain, *p <0.0001*). Error bars are 95% CI. E) Coefficient of variation of cell width of wider mutants and wild type. F) Scatter plot of the coefficient of variation of cell width versus average cell width of the entire fixed cell library. Orange dots represent the round mutants and black dots represent WT cells. r = Pearson’s coefficient.

Similarly, 40 mutants distributed over six bins (grey) (**Fig. 2A**) have a high foldchange indicating they may be wider than WT cells. These bins include the round mutants with higher IDDs that we are excluding from this analysis (**Fig. 1A**). Live cell imaging was performed on the 12 remaining strains from these upper bins at an early exponential phase (OD 0.2-0.25) and all were found to be wider than WT (*p* <0.0001) in agreement with the initial screen (**Fig. 2D**). Therefore, when including the round mutants, all 40 strains are wider than WT suggesting that the initial screen can accurately measure larger cell widths. We measured the coefficient of variation of width and found that these 12 wider cells have a larger coefficient (**Fig. 2E**) suggesting that the width of large cells varies more across the population. Since both the thinnest and fattest cells seem to correlate with the coefficient of variation of width, we measured the correlation of C_width_ to width across the entire fixed cell library. In contrast to previous results, this analysis found a high correlation suggesting that as cell width increases there is more variation in the width across the population (Pearson’s *r* = 0.8290) (**Fig. 2F**).

We performed a second analysis on the cell width by calculating the width foldchange against the plate-wise average cell width to account for intraplate variation. Among the 63 thinnest mutants comprising the black bins (**Fig. S4**) in this new analysis, 38 overlapped with the previously calculated thin group (**Fig. 2, S3**). We quantified the width of cells after live cell imaging for the four mutants with the smallest ratio out of the 25 remaining strains unique to this analysis. These four show a significant decrease of 0.89% to 18.11% (*p* <0.0001, *p* <0.05 (V187A)) of cell width than WT cells (**Fig. S3B**). Similarly, 60 mutants in the six dark grey bins (**Fig. S3A**) have a large ratio, suggesting they may be wider than the plate average. After excluding the spherical strains, Δ*rodZ,* and overlap from the initial analysis, we are left with 16 new strains. We imaged nine of these mutants and found that all of them were significantly wider (*p* <0.0001) than WT cells showing a total increase of 8.06% to 40.38% increase in cell width (**Fig. S3D**). Between these two methods we have shown that large-scale imaging of fixed cells accurately represents changes in cell width in both directions.

### MreB mutations result more in shorter cells

The activity of the elongasome adds cell length during the pre-division stage of the cell cycle. The length of *E. coli* has been linked to division time and hence the activity of the division machinery (divisome)^74,75^. It is unclear how MreB interacts with the divisome, although in *C. crescentus* MreB has been shown to condense to the division plane and in *E. coli* some labs have reported an increase of MreB at the division plane^76^. In *E. coli*, RodZ localizes at division sites in an FtsZ-dependent manner and RodZ also interacts with MreB; however, many other labs fail to see MreB localization at the division site in *E. coli*^26,33,76^. Additionally, MreB can directly interact with FtsZ which is needed to shift from elongation to division in some bacteria^38,77–79^. Hence, there may be transient crosstalk between these two complexes. If MreB is involved in regulating division, one would expect to see mutants that are affected for cell length and possibly filamentous cells in our library.

As we did for cell width, we measured the cell length foldchange of the mutants to the WT from each plate and the plate-wise average and presented the data as histograms (**Fig. 3**). The first analysis yields 65 mutants (black bar, Fig. 3A) that have a smaller ratio than WT cells suggesting they may be shorter than WT. We quantified the cell length of the 27 mutants with the smallest ratio from this group using live cell imaging and found 21 (77.78%) show significantly decreased cell length by 5.26% to 28.25% (*p* <0.0001) with most of the strains showing a smaller CV_length_ than WT (**Fig. 3B-C**), suggesting these point mutants have more control over their cell length than WT. Similarly, 16 mutants (dark grey) have a high foldchange indicating they may be longer than WT. 7 of these were excluded from future analysis because they are spherical (**Fig. 1**) leaving only 9 high ratio mutants. To further test our accuracy on measuring long cells, we looked at the next 7 rod shaped strains with the largest foldchange. Of these 16 mutants, 10 (62.5%) were significantly longer showing 3.06% to 39.80% increase in average cell length with a higher CV_length_ than WT after live cell imaging **(Fig. 3D-E**), although none formed filamentous cells.

**Figure 3:**
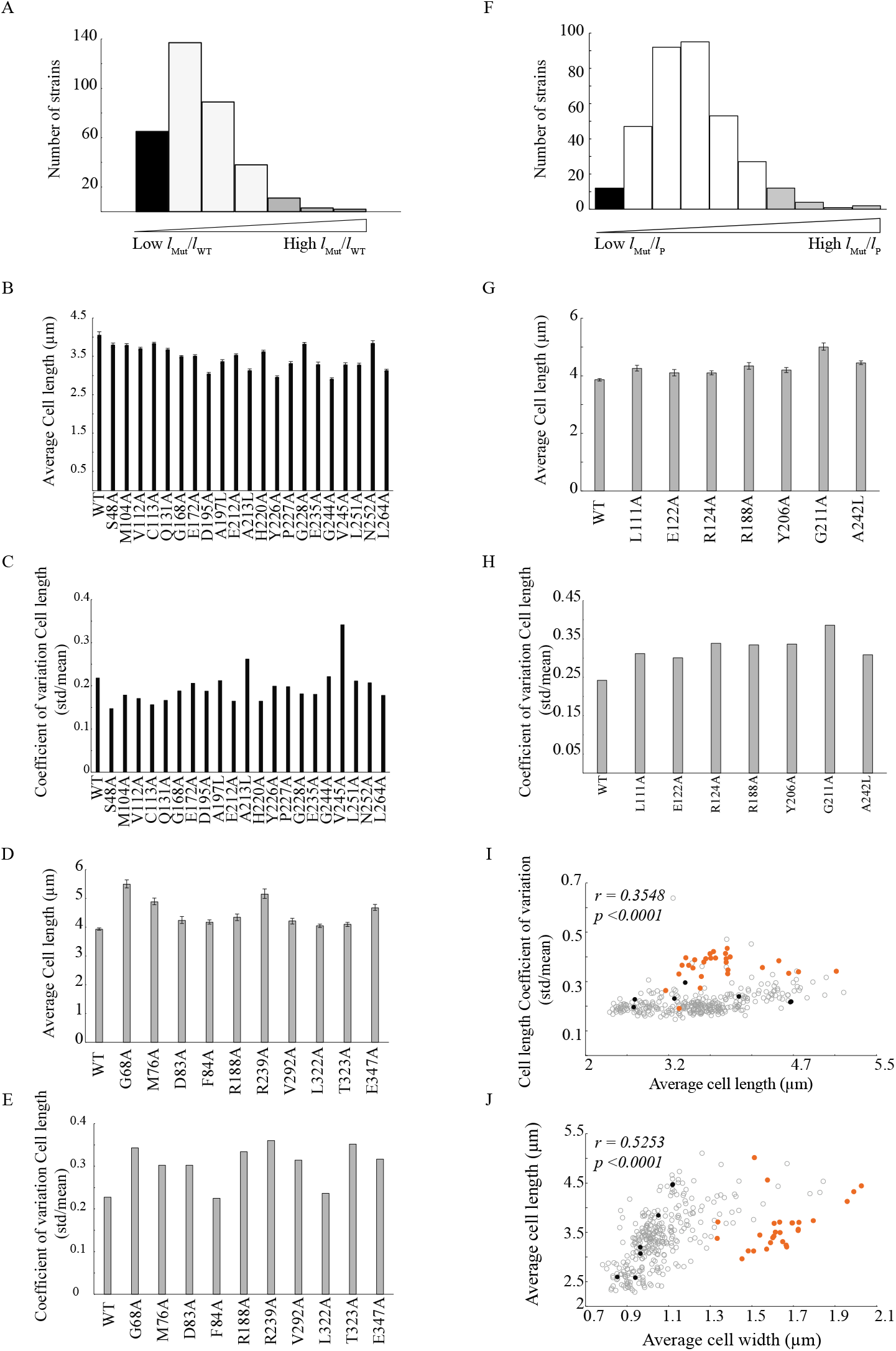
MreB has a slight effect on cell length. A) Histogram representing the cell length foldchange of 337 mutants (n >100 cells for each strain) compared to WT based on the WT cells on each 96-well plate. The black bar represents the shorter mutants, and the gray bars represent wider mutants than WT cells. B-E) Live cell imaging of cells grown in LB at 37°C to an OD 0.2-0.25. Data is pooled from three independent experiments. Error bars are 95% CI. B) Average cell width of twenty-one short mutants and WT (n >400 cells for each strain, *p <0.0001*). C) Coefficient of variation of cell width among the short mutants and WT. D) Average cell length of nine longer mutants and WT (n >400 cells for each strain, *p <0.0001*). E) Coefficient of variation of cell length of wider mutants and WT. F) Histogram representing cell width foldchange of each mutant compared to the plate-wise average length. The black bar represents the shorter mutants, and the gray bars represent wider mutants. G-H) Live cell imaging of cells grown in LB at 37°C to an OD 0.2-0.25. Data is pooled from three independent experiments. G) Average cell width of seven longer mutants and WT (n >400 cells for each strain, *p <0.0001*). Error bars are 95% CI. H) Coefficient of variation of cell length of longer mutants and WT. I) Scatter plot of cell length coefficient of variation versus average cell length from the fixed cell library. J) Scatter plot average cell length versus average cell width from the fixed cell library. I-J) Orange dots represent the round mutants and black dots represent WT cells. r = Pearson’s coefficient.

We calculated the foldchange against the plate-wise average to account for the intraplate variances. Among the 12 shortest mutants (black bar **Fig. 3F**), 11 overlapped with the previous short mutant list, leaving only one mutant; therefore, we didn’t examine short mutants from this analysis. Among the 19 longest mutants from the four bins (dark grey, **Fig. 3F**), 6 were round mutants and 2 overlapped with the list from the previous analysis, leaving one WT and 10 possibly long mutants. Among the ten, 7 (70%) mutants were significantly longer showing 6.20% to 29.52% increase in average cell length with higher variability than WT (**Fig. 3G-H**) after live cell imaging. Collectively, these results suggest longer mutants have less control on regulating cell length. We measured the correlation between the cell length and the coefficient of variation of length in the entire fixed cell library, revealing only a weak correlation (**Fig. 3I**). While the initial large-scale screen and live imaging both accurately report on cell length, it is not as robust as measuring cell width (**Fig. S5**). Similarly, there was only a mild correlation between cell length and width where we did not observe long cells within thin mutants (**Fig. 3J**). Additionally, the fact that we did not observe filamentous cells at all suggests any effects of MreB on FtsZ function is minimal.

### MreB is absolutely necessary for rod shape

When the main proteins of the elongasome are inhibited, cells become spherical, suggesting each of these proteins is important for rod shape. Previous studies using the spherical *rodZ* mutant identified suppressor mutants in *mreB* and other elongasome proteins that rescue rod shape, indicating that RodZ is not necessary for rod shape^24,25,27^. Due to the essential nature of MreB it has been difficult to do similar experiments to test if MreB is necessary for rod shape. We found 27 mutants that stably grow as spheres allowing us to test the hypothesis that unlike RodZ, MreB is required for rod shape.

To determine if MreB is in fact necessary for rod shape determination we performed a suppressor screen on six of the spherical mutants with different MreB fluorescent patterns (**Fig. 1D** and **Table 1**) to screen for different MreB defects. We used sublethal concentration of the PBP3 targeting antibiotic cephalexin to select for rod-shape suppressors. We found rod shaped mutants for four of the round-shaped parents; I15A, M90A, P108A and V181A (**Table 1**). We sequenced the *mreB* from a selection of the rod-shaped mutants and found that almost all strains have reversions back to the WT *mreB* sequence. The two mutants of the P108A spherical parent have intragenic mutation in *mreB*: F84V and G168A (**Fig. 4**). To further confirm that these mutations are the cause of the restoration of rod shape we performed WGS on these two strains and did not find any additional mutations (**Table 1**). Transduction of the *mreB* double mutants into a clean WT background results in a rod-shaped cell (data not shown). Together these results suggest the suppression comes from this second *mreB* mutation and that MreB is absolutely required for rod shape.

**Figure 4:**
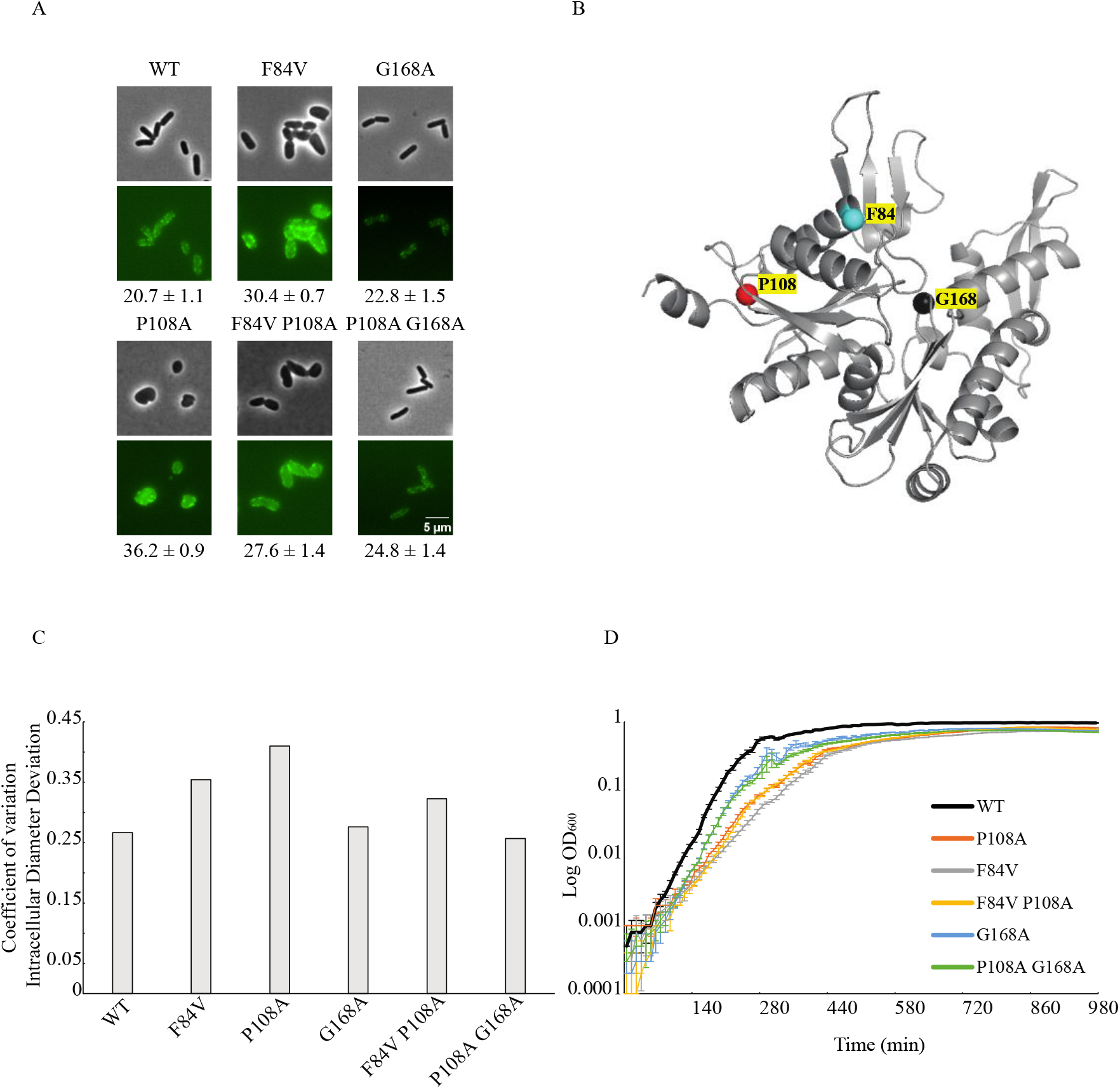
Only intragenic mutations in MreB can suppresses the round shape phenotype. A) Representative phase contrast and fluorescence images from live cell imaging of suppressor mutants, WT, and parental single mutant strains grown to early exponential phase. The numbers represent the average max doubling time (mean (min) ± sd). B) The parent mutant P108A and suppressor sites F84V and G168A are mapped on an *E. coli* MreB homology model. P108A and F84V are predicted to be in subdomain IA while G168A maps to subdomain IIA. C) IDD measurements of WT and mutants (n >700 cells for each strain). Cells were grown in LB at 37°C. D) Representative growth curves of WT (black line), suppressors, and parental strains. Cells were grown overnight in LB and sub-cultured in equal numbers into a 96-well plate with ten replicates for each strain. Error bars are standard deviations from those 10 replicates.

**Table 1:**
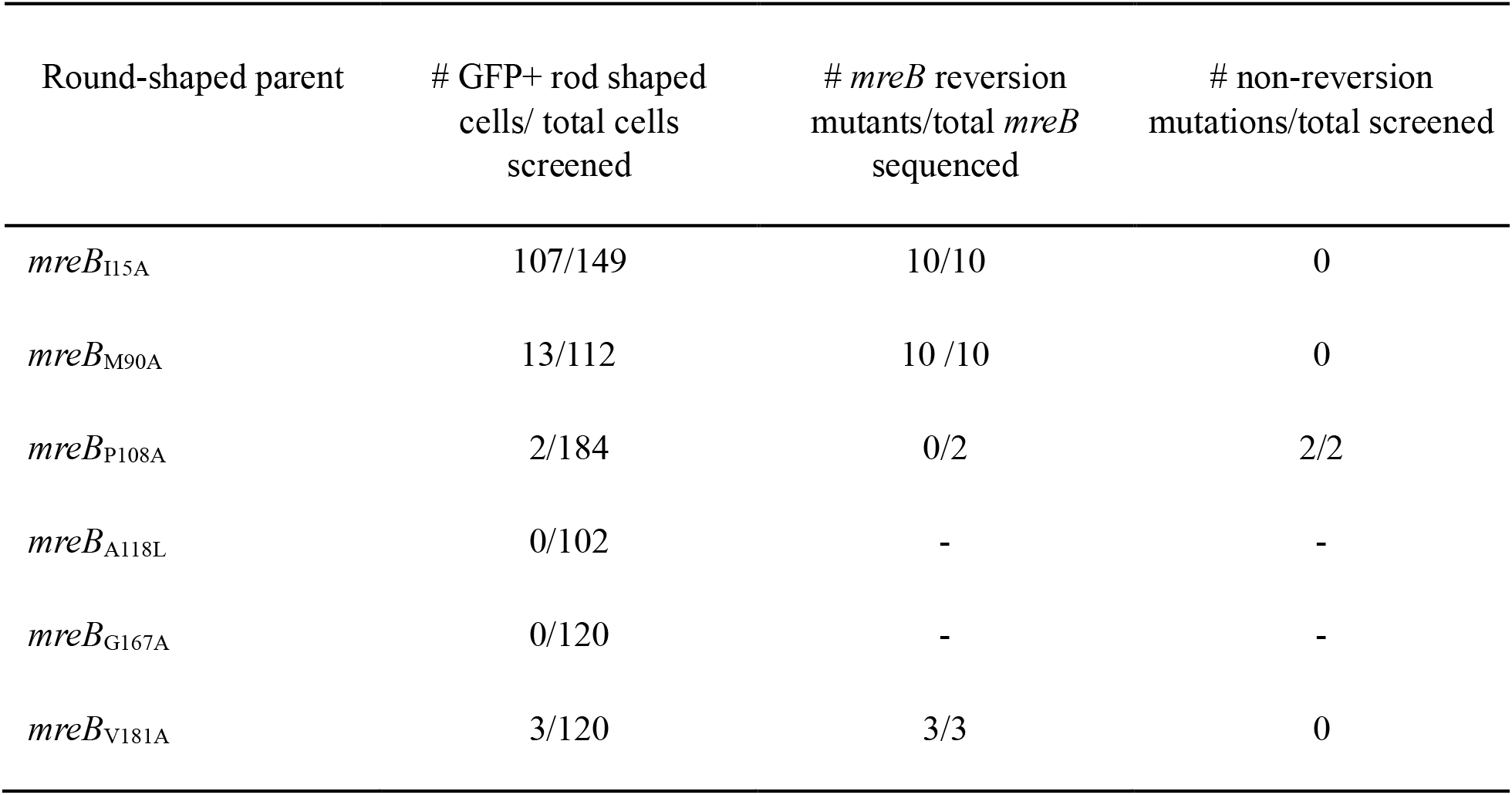
Suppressor screen of round mutants with sub-lethal level of cephalexin.

| Round-shaped parent | # GFP+ rod shaped<br>cells/ total cells<br>screened | # *mreB* reversion<br>mutants/total *mreB*<br>sequenced | # non-reversion<br>mutations/total screened |
| --- | --- | --- | --- |
| *mreB*<sub>I15A</sub> | 107/149 | 10/10 | 0 |
| *mreB*<sub>M90A</sub> | 13/112 | 10 /10 | 0 |
| *mreB*<sub>P108A</sub> | 2/184 | 0/2 | 2/2 |
| *mreB*<sub>A118L</sub> | 0/102 | - | - |
| *mreB*<sub>G167A</sub> | 0/120 | - | - |
| *mreB*<sub>V181A</sub> | 3/120 | 3/3 | 0 |

To quantify the suppressive effects of the F84V and G168A mutations we measured the IDD of both single and double mutants compared to WT (**Fig. 4C**). We found that as previously reported F84V forms a misshapen rod, while G168A has minimal effect on rod shape^46^. Both double mutant strains show a decrease in IDD, but G168A restores the rod shape to P108A better than F84V most likely due to the shape of the G168A parent being more rod-like than F184V (**Fig. 4A**). As the spherical P108A was shown to have a growth defect (**Fig. 1D**), we also measured the growth rate of the suppressors and found that they have faster doubling times than parent P108A (**Fig. 4D**). Collectively, all these observations suggest *mreB* is central in determining and maintaining rod-shape and cannot be bypassed.

### Spherical *mreB* mutants no longer use the elongasome

Round *E. coli,* such as when *rodZ* is deleted or our spherical mutants, must have a cell wall, as wall less cells (L-forms) would not be able to grow in these conditions. It is unclear how spherical *E. coli* builds their wall. A22 treatment is thought to depolymerize MreB thereby inactivating the rod system. Δ*rodZ* cells have been hypothesized to be round due to the mislocalization of MreB and therefore the elongasome, while A22 treated cells are thought to lack elongasome activity all together. As we can stably grow spherical *mreB* mutants, we hypothesize that they must be producing a cell wall in an MreB-independent manner. To confirm MreB is no longer active in these round mutants we grew the 18 strains with the highest IDD (**Fig. 1**) with and without A22 (10µg/mL) for six hours and determined the growth ratio between A22 and LB only conditions. All round mutants were able to grow better than the WT strain in A22 indicating that even in the absence of A22 these mutants do not have functional MreB to organize and coordinate the elongasome complex (**Fig. 5A**).

**Figure 5:**
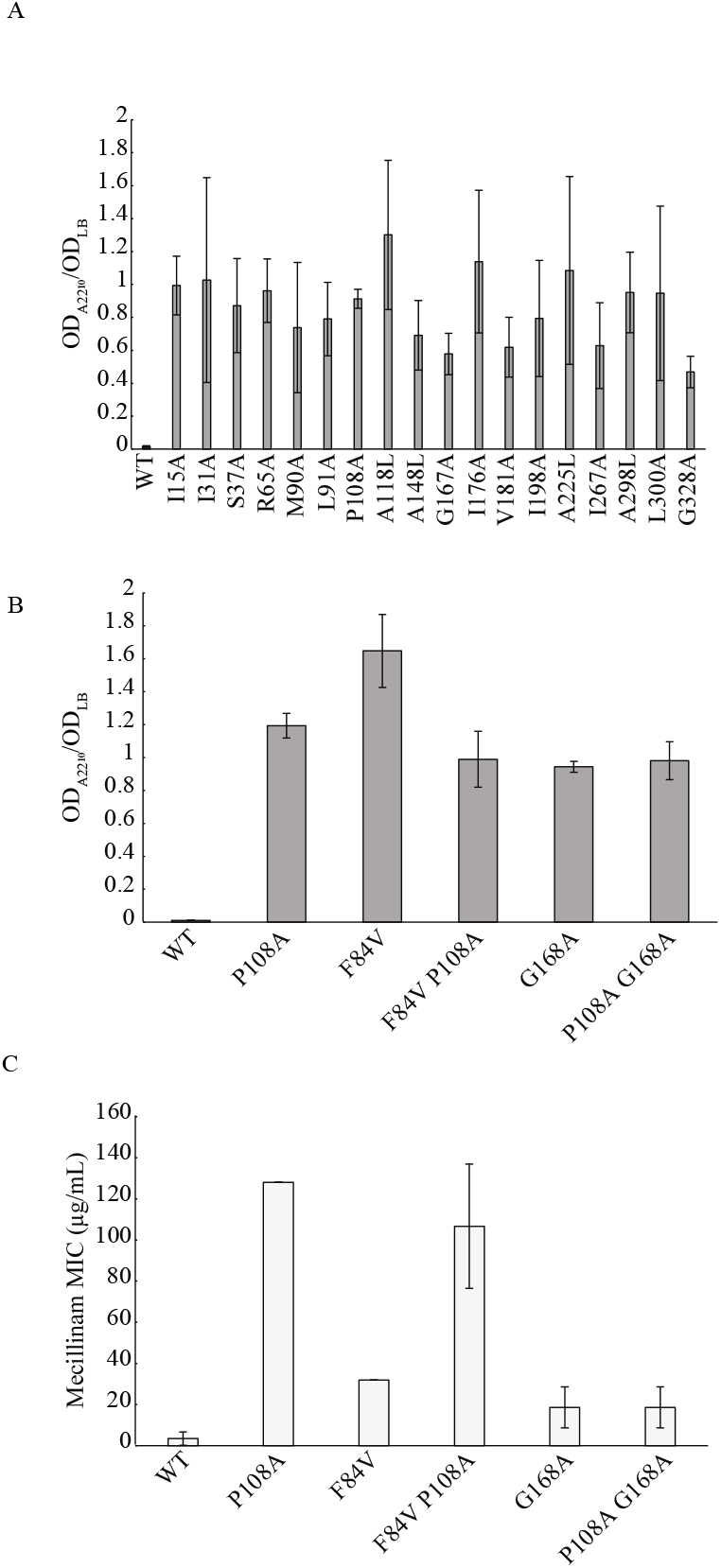
Round mutants do not require elongasome activity for PG synthesis. A-B) Cells were grown for six hours in LB or LB + A22 (10 µg/mL). The ratio of the OD between the two conditions was determined as a growth ratio. A) Growth ratios of the 18 mutants with the most extreme cell shape defects. B) Growth ratios of the rod shape suppressors and parental strains. C) Minimum inhibitory concentration to mecillinam of the suppressor mutants and parental strains. All data shown are the average values from three independent experiments with standard deviation.

The A22 resistance of the spherical mutants suggests the elongasome is no longer functioning. To further test this we examined the MIC of these mutants to mecillinam, which targets the elongasome component PBP2. All round mutants are resistant to mecillinam by > 30-fold over WT cells (**Table 2**) clearly suggesting the elongasome is no longer necessary in these strains. This could be explained if a different PG synthesis machinery is being used in these strains. The elongasome complexes account for most of the synthesis of new PG strands during cylindrical growth; however, there are other components like the aPBPs, and the divisome that can contribute to the upkeep of the PG layer. To determine if other PBPs substitute for the elongasome we measured their MICs against the PBP-targeting antibiotics: cefsulodin (aPBPs), ampicillin (multiple PBPs), and cephalexin (PBP3). All round mutants are more susceptible to cefsulodin and cephalexin, except for I15A, while all are more sensitive to ampicillin compared to WT (**Table 2**). This implies the other PBPs may have a greater role in PG synthesis independent of MreB or the elongasome in these spherical mutants. It is possible that FtsZ is driving this synthesis as these mutants are sensitive to the inhibition of PBP3 which is part of the division machinery; however, there may be a novel mechanism for PG upkeep that is independent of both MreB and FtsZ.

**Table 2:** MIC of round mutants against cell wall targeting antibiotics.

|  | MIC (µg/mL) |  |  |  |
| --- | --- | --- | --- | --- |
|  | Mecillinam | Cefsulodin | Ampicillin | Cephalexin |
| **WT** | 4.67 ± 2.49 | 32 | 2.67 ± 0.94 | 18.67 ± 9.97 |
| **I15A** | 256 | 32 | 1.67 ± 0.47 | 18.67 ± 9.97 |
| **I31A** | 256 | 16 | 0.67 ± 0.23 | 5.33 ± 1.88 |
| **S37A** | 213.33 ± 60.33 | 13.33 ± 3.77 | 0.5 ± 0.35 | 3.33 ± 0.94 |
| **R65A** | 213.33 ± 60.33 | 13.33 ± 3.77 | 0.5 ± 0.35 | 4.67 ± 2.49 |
| **M90A** | 256 | 13.33 ± 3.77 | 1 ± 0.70 | 6.67 ± 1.88 |
| **L91A** | 256 | 10.67 ± 3.77 | 0.83 ± 0.82 | 3.33 ± 0.94 |
| **P108A** | 256 | 10.67 ± 3.77 | 0.91 ± 0.77 | 3.33 ± 0.94 |
| **A118L** | 213.33 ± 60.33 | 8 | 0.83 ± 0.82 | 3.33 ± 0.94 |
| **A148L** | 256 | 13.33 ± 3.77 | 1 ± 0.70 | 13.33 ± 3.77 |
| **G167A** | 256 | 8 | 1 ± 0.70 | 5.33 ± 1.88 |
| **I176A** | 256 | 32 | 2 ± 1.41 | 13.33 ± 3.77 |
| **V181A** | 213.33 ± 60.33 | 8 | 0.83 ± 0.82 | 3.33 ± 0.94 |
| **I198A** | 256 | 8 | 0.97 ± 0.77 | 4.67 ± 2.49 |
| **A225L** | 256 | 16 | 1.67 ± 1.64 | 8 ± 5.65 |
| **I267A** | 256 | 10.67 ± 3.77 | 0.97 ± 0.77 | 9.33 ± 4.98 |
| **A298L** | 256 | 16 ± 11.31 | 0.83 ± 0.82 | 4.67 ± 2.49 |
| **L300A** | 213.33 ± 60.33 | 8 | 0.83 ± 0.82 | 3.33 ± 0.94 |
| **G328A** | 256 | 13.33 ± 3.77 | 1 ± 0.70 | 6.67 ± 1.88 |

## Discussion

MreB is a conditionally essential bacterial actin homolog responsible for rod shape determination. Rod shape can be defined by multiple parameters, such as aspect ratio and cylindrical uniformity, which have been shown to be independently regulated by MreB^33^. *In-vitro* molecular-level visualization of *Ec*MreB is not feasible; therefore, high quality *in-vivo* studies are required. To this end we made an alanine-scanning native-site MreB mutant library to use genetics to provide insights on the influence of each residue of MreB on cell shape regulation. To the best of our knowledge, this is the first truly systematic study of MreB. As previously reported, we have demonstrated that specific MreB residue can independently tune cell shape. In addition, we discovered amino acid changes that allow cells to grow as spheres in conditions not permissive for growth of an *mreB* deletion, suggesting that MreB has distinct roles in rod shape determination and viability. In support of these two distinct roles for MreB, some residues appear to be critical for viability as we were unable to make nine mutations (**Table S1**). The identification of residues essential for rod shape but not viability alone shows MreB’s role for shape regulation and viability can be separated and investigated (**Fig. 1**). Here we use the spherical mutants to show that *mreB* is absolutely required for rod shape determination and cannot be bypassed by additional mutations, unlike a *rodZ* deletion in which rod shape is restored by mutations in many elongasome proteins, and that the elongasome is no longer functioning in these cells^24,27^.

### MreB is an actin homolog

The actin superfamily is defined by conserved structural motifs needed for ATP binding/hydrolysis and polymerization. MreB has all of the structural features one would expect in an actin, including the ability to bind ATP with the nucleotide binding pocket or cleft between domains I and II^28,41,71,80,81^.

The different nucleotide states modulate actin structural transitions by affecting filament dynamics and have been shown to be important in MreB polymerization in molecular dynamic simulatins^82^. Mutations in the ATP hydrolytic or phosphate interacting motifs of actin lead to cells with poor or no growth^29,83–85^. Mutations in the ATP binding pocket in both *B. subtilis* and *C. crescentus* affect cell shape^57,58^. Some of the spherical mutants we observed are in or around this ATP binding cleft or pocket. For instance, the I166A and G167A mutations are in a conserved phosphate binding loop and lead to a round phenotype, while A298L and L300A are in the same cleft and result in similar shape defects. Taken together this indicates the significance of the proposed ATP-binding cleft in affecting nucleotide activity and therefore cell shape.

Similarly, the release of inorganic phosphate is a crucial event for actin filament polymerization. In actin, the phosphate escape channel (N111-R177) is gated by asparagine and arginine residues. An N111S mutation causes rapid release of P_i_ possibly due to absence of steric hindrance by the serine substitution^86^. Interestingly, in *Ec*MreB the structural equivalent to N111 is a A118, suggesting a lack of gating. We observed that an A118L mutation in results in a round shape-phenotype with a slow doubling time. If *Ec*MreB has a similar P_i_ escape channel as actin the substitution to leucine may introduce steric hindrance that affects P_i_ release.

Cations associate with ATP and influence actin filament polymerization and stability^87^. In yeast, mutating some of these charged residues in combination was shown to be lethal, again suggesting that single mutants might not be sufficient to explore important residues^84^. In multiple bacterial species, it has been shown that MreB assembly and stability respond to cation (Ca^++^ and Mg^++^; Na^+^ and K^+^) concentration *in vitro*^31,88–91^. Mutations in these proposed *Ec*MreB cation binding residues have minimal effect on cell shape, suggesting these MreB variants retain elongasome function. Overall, our results indicate that there may be structural differences between members of the actin superfamily that are only revealed upon genetic perturbations. Additionally, there is a need to test predicted residues with multiple amino acid changes or to make multiple mutations in important domains to fully examine the importance of specific residues.

### Predicted residues for MreB interactions are not important for cell shape regulation

In our growth conditions an *mreB* deletion would be lethal; therefore, we expected to observe lethality for residues critical for MreB assembly and function. Some residues may be essential for monomer-monomer or double protofilament interactions^92^. As MreB forms antiparallel filaments it would have been expected that mutations at the predicted monomer interfaces or at the protofilament interfaces would have a large effect on cell shape or growth. Unlike a previous study where a V121E (protofilament interface) and S284D (monomer-monomer interface) mutations create extreme shape defects hypothesized to be due to defects in MreB polymerization^28^, the corresponding alanine substitution in our library maintain rod shape. In fact, most of the mutants on predicted interface interactions are rod shaped. This previous study expressed the mutant *mreB* from a plasmid in a strain constitutively expressing *sdiA.* The fact that we don’t see large cell shape changes from most of the mutations in these or other predicted residues could be due to the nature of the alanine itself, the fact that our mutants are single copy chromosomal insertions, or because multiple residues are involved in such interfaces and the effect of a single substitution is minimal.

### *mreB* point mutations can separate its viability and shape control functions

Out of nine residues that could not be mutated, a previous study has reported different amino acid substitution(s) for seven: G134, L293, L303, P304, A332, L307 and M335^56^, indicating that while alanine substitution may not be viable, other classes of amino acids are. In total, these results suggest at least two residues, T119 and E281, may be critical for viability while the specific amino acid change is vital for MreB function in others.

Some point mutants appear to cause cells to significantly lose rod shape, while still being able to grow. These residues must be important for MreBs function in regulating rod shape, although the exact role that each residue plays is still unknown. The fact that these mutants can lose rod shape but still grow suggests that MreB independently regulates rod shape and viability.

In previous studies it was shown elevated FtsZAQ levels can suppress lethality of spherical *E. coli* cells in an *mre*^−^ background without restoring rod shape ^50,51^. The viability of our round mutants without any known suppressor mutations has the equivalent effect of FtsZ overexpression, stable round shape growth. This indicates these mutants have retained the viability function but lost the rod shape regulation function. Taken together, this suggests the two functions of MreB of cell shape regulation and viability can be separated.

### Intragenic *mreB* mutations suppress the round shape-phenotype

Previously, elongasome components were shown to restore rod shape in a *rodZ* mutant^24,27^. While it has been suggested that MreB is necessary for rod shape, similar suppressor screens have been unfeasible until now. To determine if MreB can be bypassed in a similar manner to RodZ we attempted to find suppressors of the cell shape defect in six of the spherical mutants **(Table 1)**. While most of the mutations we found were reversions back to WT, we did identify two rod-shape suppressors in the P108A parent. Both suppressors are intragenic to *mreB* (**Fig. 4**). To confirm these intragenic mutations are the cause of suppression we performed WGS on the double mutants and moved these mutations into a fresh WT background via transduction. In support of the fact that these intragenic mutations are the cause of suppression there were no additional mutations in genes involved in PG synthesis or regulators in the double mutant when compared to the P108A parents and the transductants remain rod shaped (data not shown). G168A is a better suppressor of the round shape than F84V (**Fig. 4C**) and is in the ATP binding pocket. It may influence ATP binding/hydrolysis possibly leading to a more stable polymers^92^. In support of this, the G168A mutation results in A22 resistance (**Fig. 5**). The suppressor strain with the P108AG168A mutation is much more sensitive to mecillinam than the P108A parent, suggesting it may have restored the elongasome activity that was lost in the P108A parent. Overall, the results suggest the shape maintaining function of MreB cannot be compensated by other elongasome components making MreB essential for rod-shape maintenance.

### Round mutants’ PG synthesis may be independent of elongasome activity

The activity of PBP2, a transpeptidase (TP) that crosslinks muropeptide strands is crucial for elongasome activity. The loss of rod shape in our round mutants suggests that the activity of the elongasome is perturbed. To test this, we measured the susceptibility of 18 round mutants to mecillinam, an amidinopenicillin that specifically inhibits the TP activity of PBP2. All the mutants tested were resistant (>30 folds than WT), suggesting that PBP2 activity is no longer required and therefore that the elongasome is not functioning in these mutants (**Table 2**). In addition, we measured the MIC of these mutants against cefsulodin (targets: PBP1A and PBP1B) and cephalexin (targets PBP3). All mutants but one (I15A) were more sensitive to them indicating other major PBPs may have a greater role in PG synthesis in these strains (**Table 2**). PBP3 is part of the divisome suggesting that there may be increased divisome activity in these mutants and FtsZ has been shown to direct PG incorporation into the lateral cell wall when MreB is perturbed^93^. This suggests that the suppressive nature of overexpression of FtsZAQ is not due to making a larger Z-ring as previously suggested, but rather to increasing general cell wall synthesis allowing the cells to grow without a functioning elongasome.

## Materials and Methods

### Bacterial growth

Bacteria were grown using standard laboratory conditions. Unless stated otherwise, cultures were grown overnight in LB medium (10 g/L NaCl, 10 g/L tryptone, 5 g/L yeast extract) and subcultured in the morning 1:1000 at 37°C in a shaking incubator.

### Library construction

An ordered array of alanine or lysine substitutions was made by Twist Bioscience consisting of a linear DNA molecule including DNA upstream of *mreB,* a kanamycin resistance cassette, and the *mreBCD* operon (**Fig S1A**). Each construct has a *gfp*-sandwich fusion between a non-conserved loop of *mreB*^46^. The constructs were amplified using PCR and transformed into an *E. coli* MG1655 strain with the PKD46 plasmid that facilitates λ-red recombination^94^. The recombinants were selected on LB plates supplemented with kanamycin (30 μg/mL) at 30°C and screened for GFP fluorescence. Suspension of desired colonies were incubated at 42°C to cure them of PKD46 plasmid. Recombinants with mutations near the C-terminal region of *mreB* were sequence verified to ensure proper recombination occurred.

### Library fixation

Strains were sub-cultured in 96-well plates from freezer stocks in LB overnight at 37°C. Overnight cultures were diluted 1:1000 in fresh medium and grown for three hours. Cultures were 1:1000 diluted into fresh medium and grown for two more hours and then diluted 1:1000 a second time to keep the in exponential phase and grew for two hours. Cells were fixed with paraformaldehyde for 15 minutes, washed with PBS three times and resuspended in PBS.

### Microscopy

For initial imaging of the library see Library Fixation for cell prep. For all live cell imaging, cultures were diluted to equal numbers (0.1 OD) and 2 μL of suspension was inoculated into 2 mL of LB and incubated at 37°C to early exponential phase (0.2 to 0.25 OD). All analysis from live cells includes data pooled from three independent experiments.

All imaging was done on 1% M63-glucose agarose pads at room temperature. Phase contrast and fluorescent images were collected on a Nikon Ni-E epifluorescent microscope equipped with a 100X/1.45 NA objective (Nikon), Zyla 4.2 plus cooled sCMOS camera (Andor), and NIS Elements software (Nikon).

Cell shape statistics were calculated using the MATLAB software Morphometrics^56^, and custom software as described previously^25^. Only single non-diving cells were used for analysis unless stated otherwise. For a list of data from the fixed cell imaging see supplemental data file 1. Pearson correlation coefficient was used to analyze relationships between sets and Student’s t test was used for statistical analyses. All data from the fixed cell imaging can be found in supplemental data file.

### Growth rate analyses

Overnight cultures were grown in LB at 37°C and diluted in LB in equal numbers (0.1 OD) as described previously. 1 μL of the suspension was inoculated into 100 μL of LB in 96-well plates for each strain with at least three biological replicates on each plate and repeated three times. Plates were incubated at 37°C shaking for 16hr in a Biotek plate reader. Doubling times are reported as the max doubling time as determined by the maximal instantaneous growth rate.

### Suppressor screen analysis

Overnight cultures were grown in LB at 37°C and diluted in LB in equal numbers (0.1 OD) as described previously and grown to 0.3 OD. 100 μLs of cells were plated onto LB plates supplemented with differing concentrations of cephalexin (3.0, 4.0, 5.0, 6.0 & 7.0 μg/mL) in triplicate and incubated overnight at 37°C. Isolated colonies were suspended in 10 μL LB and imaged to analyze the cell shape.

### Sequencing

For whole-genome sequencing, genomic DNA was extracted from strains from overnight cultures using Qiagen DNeasy Blood and tissue kit. Illumina sequencing was performed by MIGS (now SeqCenter) and variant analysis was performed using Breseq^95^.

For single construct sequencing, the entire *mreB* cloning construct or *spoT* were PCR amplified and sequenced using nanopore technology (Plasmidsaurus).

### Mapping mutations

Pymol was used to map the mutations on the AlphaFold model of *Ec*MreB.

### Antibiotic sensitivity profiling

Overnight cultures were diluted into equal numbers (0.1 OD) and 1 μL was inoculated into 100 μL of LB supplemented with a gradient of antibiotic concentrations of two-fold dilutions. Mecillinam (512 μg/mL), Cephalexin (16 μg/mL), Cefsulodin (16 μg/mL) and Ampicillin (16 μg/mL). Plates were incubated at 37°C for 16h with shaking. The MIC was defined as the concentration of drug that resulted in an OD less than 0.1 times the OD of the control well without any drug. For six-hour growth ratio assay, overnight cultures were inoculated in equal numbers into 2mL of LB with and without with different antibiotic concentrations (For A22, 1.0 and 10μg/mL) and incubated at 37°C for 6h with shaking. It was repeated for three separate days.

